# Neuronal and Astrocyte Insulin-like Growth Factor-1 Signaling Differentially Modulates Ischemic Stroke Damage

**DOI:** 10.1101/2023.04.02.535245

**Authors:** Cellas A. Hayes, Nyah I. Morgan, Kamryn C. Thomas, M. Jake. Pushie, Akshaya Vijayasankar, Brandon G. Ashmore, Kendall Wontor, Miguel A. De Leon, Nicole M. Ashpole

**Affiliations:** Department of BioMolecular Sciences, University of Mississippi School of Pharmacy, University of Mississippi, University, MS 386671; Department of Surgery, College of Medicine, University of Saskatchewan, SK S7N 5E5 Canada; Department of Chemistry and Biochemistry, The University of Mississippi, University, Mississippi 38677, United States; Research Institute of Pharmaceutical Sciences, University of Mississippi School of Pharmacy, University of Mississippi, University, MS 38677

**Keywords:** stroke, IGF-1, Somatomedin C, glia-neuron interaction, neurovascular unit, neuroprotection, astrocytes

## Abstract

Ischemic stroke is a leading cause of death and disability, as therapeutic options for mitigating the long-term deficits precipitated by the event remain limited. Acute administration of the neuroendocrine modulator insulin-like growth factor-1 (IGF-1) attenuates ischemic stroke damage in preclinical models, and clinical studies suggest IGF-1 can reduce the risk of stroke and improve overall outcomes. The cellular mechanism by which IGF-1 exerts this protection is poorly defined, as all cells within the neurovascular unit express the IGF-1 receptor. We hypothesize that the functional regulation of both neurons and astrocytes by IGF-1 is critical in minimizing damage in ischemic stroke. To test this, we utilized inducible astrocyte-specific or neuron-specific transgenic mouse models to selectively reduce IGF-1R in the adult mouse brain prior to photothrombotic stroke. Acute changes in blood brain barrier permeability, microglial activation, systemic inflammation, and biochemical composition of the brain were assessed 3 hours following photothrombosis, and significant protection was observed in mice deficient in neuronal and astrocytic IGF-1R. When the extent of tissue damage and sensorimotor dysfunction was assessed for 3 days following stroke, only the neurological deficit score continued to show improvements, and the extent of improvement was enhanced with additional IGF-1 supplementation. Overall, results indicate that neuronal and astrocytic IGF-1 signaling influences stroke damage but IGF-1 signaling within these individual cell types is not required for minimizing tissue damage or behavioral outcome.

## 1. Introduction

Stroke and other cardiovascular diseases continue to wreak havoc as one of the leading causes of death and disability worldwide. Over 5.5 million individuals have died from stroke, and many survivors are left with physical impairment, cognitive dysfunction, loss of independence, and financial burdens [1]. Ischemic strokes account for more than 80% of strokes that occur [2]. In general, ischemic strokes occur through the blockage of a blood vessel which causes an abrupt loss of oxygen and glucose deficiency leading to ATP depletion, the generation of oxidative stress mechanisms, excitotoxicity, necrosis, apoptosis, ferroptosis, and ultimately a brain infarct [3, 4]. Preclinical studies have identified protective interventions capable of ameliorating these drivers of neurodegeneration; however, most treatments have failed to translate to clinical populations [5-7]. However, insulin-like growth factor-1 (IGF-1) has been proposed as a preclinical and clinical solution to combating ischemic stroke.

Circulating IGF-1 levels are inversely related to stroke incidence [8-12]. Numerous clinical studies have shown that reductions in IGF-1 levels are also associated with poorer clinical outcomes following stroke [8-12]. Moreover, several preclinical rodent studies show exogenous IGF-1 reduces ischemic infarct size and accompanying neurological deficits [13-18]. Nevertheless, there is a significant gap in knowledge regarding the molecular and cellular mechanisms by which IGF-1 exerts neuroprotective effects against ischemic stroke.

IGF-1, previously referred to as Somatomedin C, is a 70 amino acid polypeptide classified as a neuroendocrine modulator with autocrine and paracrine functions. IGF-1 is primarily hepatic derived where it is produced by hepatocytes and secreted into the bloodstream to cause an array of pleiotropic effects. Within the brain, neurons and neuroglia can both produce IGF-1 locally. Whether made in the liver or by other local tissues, IGF-1 signaling is mediated by the insulin-like growth factor-1 receptor (IGF-1R) ubiquitously expression on all major cell types throughout the body, including neurons and neuroglia within the brain [19-21]. As described in our literature review [22], IGF-1 has the potential to act directly on neurons to provide neuroprotection following ischemic stroke or indirectly by modulating the functional responses of supporting glial cells within the damaged area. In general, neurons are the first to undergo cellular death events due to the lack of oxygen and ATP, while astrocytes readily buffer extracellular glutamate and begin to form the glial scar around the ischemic core. Other cells such as endothelial cells and microglia are also impacted by ischemic insults, contributing to changes in blood brain barrier permeability and increased inflammatory responses, respectively. Overall, increased IGF-1 signaling within each of the cell types in the brain could benefit from their role in combating ischemic stroke.

The current study addresses this gap in mechanistic understanding by exploring the immediate and short-term changes that occur when IGF-1 signaling is manipulated in the two major cell types of the brain, astrocytes and neurons. We hypothesized that the functional regulation of both neurons and astrocytes by IGF-1 is critical to minimize damage in ischemic stroke. To test our hypothesis, we utilized inducible transgenic models where we could selectively reduce IGF-1R in adult astrocytes or neurons and induce stroke via ischemic photothrombotic stroke (PTS). This approach allowed for a delineation of the importance of IGF-1 regulation of neurons versus IGF-1 regulation of astrocytes during ischemic stroke on the subsequent outcome. We assessed blood brain barrier damage, microglial infiltration, biochemical changes, and neurological responses in the hours following stroke.

Furthermore, we examined the effects of intranasal IGF-1 within our transgenic mice to determine whether astrocytic IGF-1R and/or neuronal IGF-1R was necessary for the protective effects of exogenous IGF-1. Together, this study provides insight into the path by which IGF-1 affords neuroprotection against ischemic stroke.

## 2. Materials and Methods

Animals: All methods and procedures were approved by the Institutional Animal Use and Care Committees of the University of Mississippi. Mice were housed in ventilated racks (Lab Products, Inc. RAIR Envirogaurd ventilated cages) with enriched bedding under a 12:12 hour photoperiod, with 5053 Pico Lab chow and water ad libitum. Male and female mice were housed in cages 3-5, and random block randomization was used to ensure equal group size for study enrollment.

Original transgenic mouse strains were purchased from Jackson laboratories and crossed in house to develop experimental cohorts. Female igfrf/f (B6;129-Igf1rtm2Arge/J) mice expressing Lox P sites surrounding exon 3 (encoding ligand-binding domain of IGF-1R) were crossed with male igfap-Cre/ERT+ mice for the astrocyte specific studies or male B6;129S6-Tg (icamk2a-cre/ERT2)1Aibs/J mice for the neuronal specific studies. F1 generation male igfrf/+ expressing the Cre transgene were bred with igfrf/f females for a second generation (F2). For the astrocytic studies, the offspring igfap-cre/ERT2-igfrf/f and igfrf/f mice (F3+ generation) were enrolled in the study; similarly, for the neuronal studies, F3+ mice expressing icamk2-Cre-igfrfl/fl and igfrf/f were enrolled. Mice were genotyped to verify igfrf/f homozygosity and igfap-Cre/ERT or icamk2a-cre/ERT2 expression using the primers and suggested PCR protocols designed by Jackson laboratories, as previously reported (Prabhu, Khan et al. 2019). Following the RED Extract-N-Amp Tissue PCR XNAT manufacturer’s protocol (Sigma), 2-3mm of each mouse’s tail was removed and DNA was extracted. The recommended primers and PCR protocols designed by Jackson laboratories were used to verify the transgenes (primers purchased from Sigma). Following PCR amplification, the products were run on a 2% agarose gel stained with green glow (Denville) at 120V in 1XTBS for 1 hour. UV imaging identified PCR products. All gels contained a 100bp DNA ladder (ThermoFisher) and a designated positive control for comparison.

IGF-1R Reduction [23]: The designated knockout mice received tamoxifen to induce Cre recombinase expression in the transgenic mice at three months of age. Per Jackson laboratory’s recommendation, all mice received tamoxifen to remove the floxed sequences of the target gene. Tamoxifen was diluted in corn oil, solubilized at overnight at 50°C, cooled, and administered via i.p. injection for 5 consecutive days in a 75mg/kg dose. It has recently been shown that tamoxifen induces neurogenesis (Smith, Saulsbery et al. 2022); therefore, control igfrf/f mice (absent of Cre gene expression) also received tamoxifen.

Vaginal Cytology [24]: To account for the influence of estradiol on neuroprotective mechanisms, female mice received a stroke during the metestrous or diestrous phase of their estrous cycle for infarct analysis cohorts. Briefly, epithelial tissue was collected using 70μl of double distilled water and assessed using light microscopy with a 10X objective. Mice were cycled two hours following the light cycle change (9am) and all surgeries were performed within a four-hour window following estrous cycle determination. The estrous cycle of all female mice was documented and did not reveal effects of metestrous vs diestrous.

Photothrombosis: Photothrombotic ischemic stroke was induced following a similar protocol as Blanco Suarez et al [25]. Male and female mice at 4-6 months of age were anesthetized with 5% isoflurane and 1.2% oxygen, placed on stereotaxic frame, and maintained under the anesthetic plane with 2% isoflurane and 1.2% oxygen via nose cone. Mice were intraperitoneally injected with rose bengal (Fisher R323-25) in saline (0.9% NaCl) at a dose of 100µg/g (10µl/g). Rose bengal was prepared fresh and protected from light prior to each day of surgery. Following injection, the surgical scape was prepped with chlorohexidine swabbing, a 2cm midline incision was performed, and the scalp was teased apart using forceps to expose bregma. Five minutes after the rose bengal injection, the right frontal skull surface was illuminated for 10 minutes with a 520nm diode laser (Thor labs) +3mm lateral to and -0.6mm posterior to bregma (stereotaxic coordinates). The skull was then washed with 0.9% saline, the incision was closed using two 4-0 nylon thread mattress sutures, and lidocaine and prilocaine cream (2.5%/2.5) was applied to the surgical wound surface. The mouse was then returned to a fresh home cage with a heating pad under 50% of the cage and allowed to recover for one hour. Control (sham) surgical mice were injected with rose bengal and subjected to the entire procedure without laser illumination. Two mice did not survive the PTS procedure due to over exposure to anesthesia. No differences in body weight were detected between experimental groups prior to or following PTS.

Exogenous IGF-1 Administration: Human recombinant IGF-1 (R&D Systems 291-G1) was reconstituted in sterile saline and 5µls (30µgSE) was administered during inspiration to the intranasal passages of mice immediately, at 24, and 48 hours following the photothrombosis procedure [26, 27]. Intranasal injections were performed without anesthesia to avoid the confounds of anesthesia-induced neuroprotection [28].

Tissue Isolation: At the end of experimentation, mice were euthanized via rapid decapitation or through exsanguination under anesthesia (5% isoflurane and 1.5% oxygen). Once unresponsive to toe pinch, mice underwent a ventral midline incision in the thoracic cavity to expose the heart. Whole blood was taken directly from the left ventricle. The superior vena cava was cut via micro dissecting scissors, and a syringe containing ice cold 1X PBS was inserted into the left ventricle to circulate the PBS peripherally. The whole blood was centrifuged at 2000g for 20 minutes to separate serum, which was immediately frozen and stored in the -80°C for analysis. Following perfusion, brains were fixed using 4% paraformaldehyde (in phosphate buffer), immersed in 30% sucrose (in phosphate buffer), embedded in optimal cutting temperature compound (OCT; Fisher 4585), and frozen in a 100% ethanol:dry ice bath. 45µM sections were obtained using a Leica CM3050 cryostat and stored in cryopreservative until immunohistochemical staining (below). In addition, some 10µM slices were gathered and directly mounted to double frosted microscope slides for Evan’s blue quantification. Others were mounted to low E slides to quantify biochemical changes using Fourier Transform Infrared Spectroscopy (both below).

TTC Stain [29, 30]: Mice were euthanized via rapid decapitation, 0.5mL of whole blood was taken, and brains were removed and placed in -20°C for 3-5 minutes. Once tissue had slightly hardened, brains were placed in a 100-µm brain matrix (Alto 69080).

Razor blades were inserted every100-µm from the anterior to the posterior brain until six slices were acquired. Each 100-µm brain slice was placed in a 2% triphenyl tetrazolium chloride (TTC; Sigma Aldrich T8877-50G) dissolved in 1X phosphate buffer saline. Tissue was incubated in TTC solution in a 37°C incubator for 30 minutes or until the tissue was visually deep red.

Following incubation, the stained brain slices were immediately washed with 1XPBS and placed in a 4% PFA solution (paraformaldehyde). Each set of slices of a single mouse wwas placed on a microscope slide and standard HEIC images were obtained and converted to jpegs for quantification using Image J software. The front and back of each slice were imaged to account for variation and area of infarcts. Infarct size throughout the slices was quantified by a blinded observer using ImageJ and the percent infarct of those slices was calculated and compared across groups using a two-way ANOVA and Tukey’s post-hoc test.

Evan’s Blue Staining: Blood-brain barrier permeability was examined 3 hours after ischemic stroke with a protocol adapted from Manenko et al 2011. In short, immediately following PT induction, each mouse received an intraperitoneal injection of 2% Evan’s blue dye (Sigma E2129-10G) dissolved in 0.9% saline solution. The dye was allowed to circulate for 3 hours before transcardial perfusion with ice cold 1X PBS (Gibco 10010-031). Post-perfusion, the whole brain was fixed and sliced. The 10µM slices were directly mounted to double frosted microscope slides for Evan’s blue fluorescent quantification. Evan’s blue naturally fluoresces at red (546nm+) wavelengths; thus, we used this novel approach to quantify its leakage in the tissue. Coronal sections were imaged with a 4X objective on a High Content Analysis Nikon Ti2-E inverted fluorescent microscope, with TRITC filters (546ex/577em) and 9×9 large imaging stitching. The signal to background output from Nikon was used as the quantitative measure. The representative images were converted to dark rainbow to emphasize the statistical results. Data were compared using a one-way ANOVA followed by Tukey’s post-hoc test to determine difference of means.

Immunohistochemistry: Brain slices (45µM) were washed three times in 1XPBS and permeabilized in PBS Tween-20 (0.10%) for 20 minutes. Antigen retrieval was performed with boiling citrate buffer. The free-floating slices were incubated in the citrate buffer at 95°C for 30 minutes, cooled for 20 minutes, washed, and blocked using 0.5% BSA for 30 minutes at room temperature. Primary antibody was added in 0.1% BSA overnight at a dilution of (1:500 IBA-1 Wako 019-19741). Secondary antibody was added at a 1:1000 (Alex Fluor 488 Goat Anti-Rabbit Invitrogen A11034) for 2 hours at room temperature. Sections were mounted using Prolong anti-fade DAPI (Invitrogen P36931) mounting medium and sealed with a coverslip. A Leica SP8 inverted confocal microscope with a white-light laser system was used to image the Iba+ cells (40X objective, 488ex/525em). Images were acquired using a z-stack (z=1µM) of two random areas (n=2) of three to four slices (n=3-4) per animal (n=3). Unless otherwise stated, confocal microscopy images are shown at maximum projection of z-stacked images, for the evaluation of microglia number, endpoints per cell (Young and Morrison 2018). Each z-stacked image set was processed and quantified with Fiji/Image software (NIH, USA). A maximum projection of the Iba1+ fluorescent staining (488ex/525em) was generated and converted to a gray image using the lookup table. The brightness/contrast was adjusted, background subtracted (pixels=500), unsharp masked (pixel radius=3 and mask weight=0.6), despeckled, and thresholded to generate a binary image. Endpoints were closed, and bright outliers (pixel radius=2/ threshold=50) were removed. Lastly, the analyze particle plugin was used to obtain microglia cell count, and subsequently, the binary images were skeletonized and the plugin analyze skeleton 3D/2D was used to obtain branch and end point voxel data. All maximum branch lengths and endpoint voxels of two with a length less than 3.079 were removed. The sum of the end point voxels and maximum branch length were calculated. In addition to plotting and statistically comparing the raw summed end point voxels and branch length, data were normalized to the number of microglia in the image. All Iba1+ data were compared using a one-way ANOVA followed by Tukey’s post-hoc test to determine difference of means.

Fourier transformed infrared resonance spectroscopy: FTIR spectroscopic imaging was performed using a Bruker’s LUMOS II FTIR Microscope equipped with a liquid nitrogen cooled 32×32 focal plane array (FPA) detector fitted with an 8x objective. The data was acquired in reflectance mode with 64 co-added scans and 8×8 binning over the 4000-750 cm-1 spectral range with a resolution of 4 cm-1. Prior to measurements, a background (blank) image from a region adjacent to the sample was captured on a section of the low E slide without OCT or tissue. Considering the reduced penumbral size of the PTS model and the time of interest (3 hours post stroke), the ipsilateral hemisphere was imaged using 11×1 fields of view for all stroke groups, 9×1 for sham groups, and 2×1 for all contralateral hemispheres. Representative images of the contralateral and ipsilateral images were obtained using a 43×7 square scan with 16 co-added scans, a resolution of 4 cm-1, and 4×4 binning. Additional data processing was performed using the data mining and spectroscopy analysis tool Quasar, following established methods [31]. Integrated peak intensities were acquired following baseline subtraction using the rubber band method to minimize inconsistencies in the baseline from the low E slides. Second derivative peak intensities were generated following vector normalization, using a Savitzky-Golay filter, with a 13 point window and second order polynomial. Due to the tissue treatment artifacts introduced from PFA treatment data was analyzed following reference values obtained from Ali, Rakib et al. (2018) [32]. FTIR peaks listed in Table 1 were used for signal averaging from the entire region of interest in Quasar for semi-quantitative analysis [32].

**Table 1.** Spectral Ranges for Quantitative Analysis (IAUC)

| Band Assignment | Observed Wave Number (cm<sup>-1</sup>) | Integrated Spectral Range (cm<sup>-1</sup>) |
| --- | --- | --- |
| Total Protein | - | 1700-1500 |
| Amide I | 1649 | 1700-1600 |
| Amide II | 1543 | 1600-1500 |
| Total Lipid | - | 3100-2800 |
| CH<sub>2</sub> sym | 2852 | 2865-2940 |
| CH<sub>2</sub> asym | 2924 | 2935-2915 |
| CH<sub>3</sub> sym | 2961 | 2965-2955 |
| <sup>a</sup>Lipid Ester C=O | 1739 | 1755-1715 |
| Olefin = CH | 3015 | 3027-3000 |

\*Abbreviations: sym symmetric stretching; asym asymmetric stretching. <sup>a</sup> is indicative of oxidative stress

Cytokine Array: Isolated serum was run on the Luminex FlexMap 3D for inflammatory cytokine/chemokine production using the ProcartaPlex plate with 26 analytes (ThermoFisher EPX260-26088-901) per manufacturer’s protocol. Data was compared between groups using a one-way ANOVA and Tukey’s posthoc test.

Neurological Deficits Score [33]: Neurological deficits were adapted from (Rousselet, Kriz et al. 2012) [33]. Mice were scored at 3 hours for short term studies and at 1, 24, 48, 72-hours post-reperfusion with a six-point scale, whereby 0=normal, 1= ipsilateral rotating with <50% contralateral rotation attempts, 2= >50% contralateral rotating with <50% rotational attempts, 3= consistent and only contralateral rotating, 4= ipsilateral rotating with contralateral twist of posterior region, 5= severe rotation and progression into barreling, or a 6= coma/moribund.

Locomotor Asymmetry: The cylinder test assesses locomotor asymmetry and evaluates limb usage by comparing the number of independent and simultaneous placements of each forelimb on the wall of a plexiglass cylinder where the mouse is placed. Each mouse was placed in a 20 cm cylinder for 5 minutes prior to stroke induction and 72 hours post-photothrombosis. Each mouse was recorded during the duration of the task. The first 20 paw placements or all the paw placements were quantified if the mouse did not reach a total of 20 during the 5-minute duration. The percent paw usage ((72hr # - Pre #) / Pre #)*100) and the paw placement preference discrimination index (72hr # - Pre #) / ((72hr # + Pre #) + Pre #)) were calculated.

Statistical Approach: Statistical analysis and graphical analyses were performed using Sigma Plot version 14 software and R studio. One-way and two-way ANOVAs were used when appropriate (defined in the figure legend). Post-hoc comparisons were selected based on the experimental question and fulfillment of normality and equal distribution. Details for each are provided in the figure legend or independent method section. No sex difference was detected in infarct size. Thus data for males and females were combined, as delineated. All data were expressed with mean +/- standard error. For all studies, p<0.05 was used as the statistical significance value and denoted using different alphabetical characters. A full table of statistical comparisons is found in the Supplementary Materials.

## 3. Results

### 3.1 Astrocytic IGFR Contributes to Neurological Response and Blood Brain Barrier Integrity in the Hours Following Stroke

Transgenic mice expressing igfap-Cre-igfrfl/fl or icamk2a-Cre-igfrfl/fl were generated to allow for Cre-remediated induction of astrocyte-specific IGFR knockout (aKO) or neuron-specific IGFR knockout (nKO), respectively (Figure 1A[23]). Cre recombination was induced with tamoxifen in adult mice (3 months of age) to reduce the confound of developmental defects.

**Figure 1.**
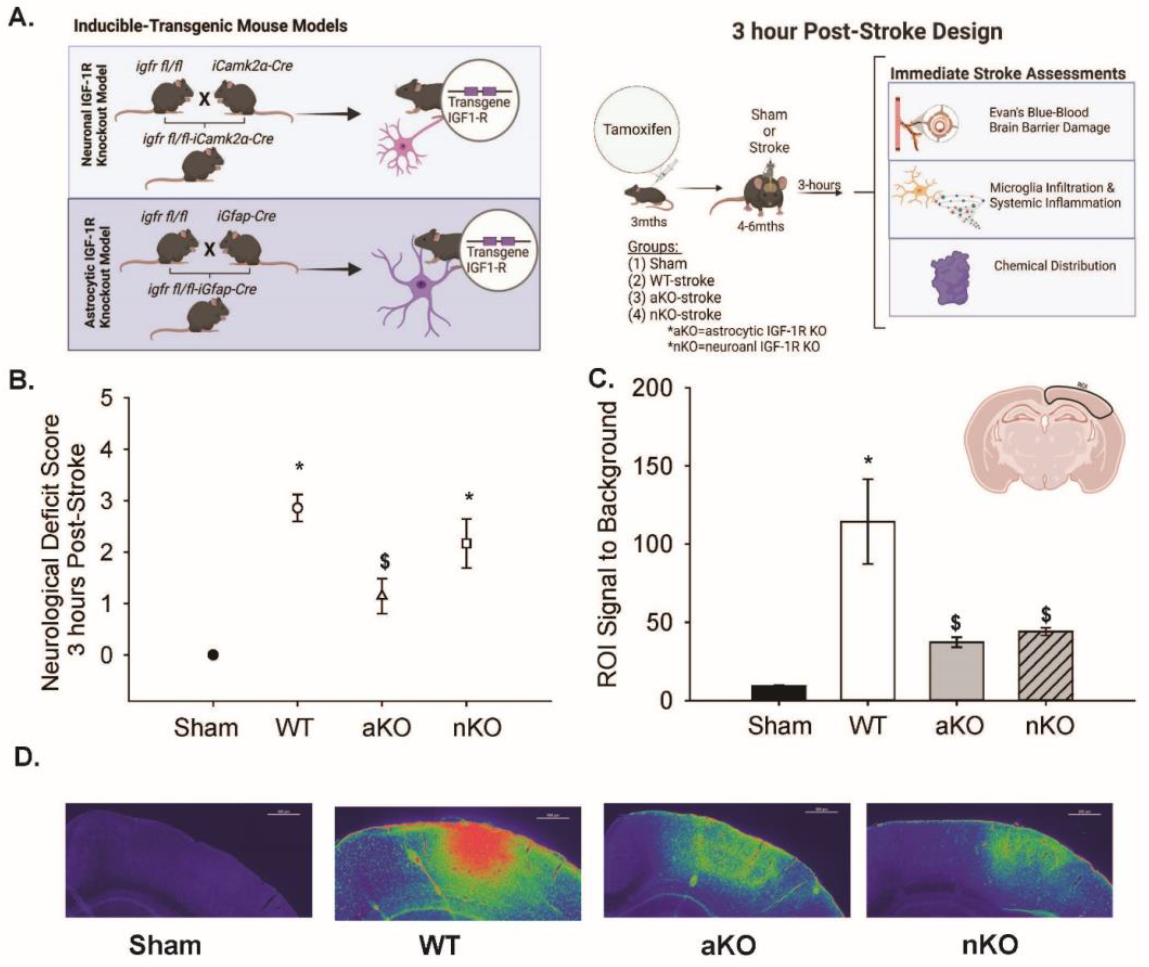
Experimental Design of Immediate Stroke Associated Changes and Blood Brain Barrier Changes. (A) Tamoxifen inducible transgenic mouse models to reduce IGF-1R in neurons or astrocytes. Design and timeline of neuron (nKO) or astrocyte-specific (aKO) deletion of IGF-1R impact on immediate stroke changes (3 hours post-insult). (C) ROI) signal to background noise between Sham, WT, aKO, and nKO groups (n=16-37). (D) Representative images of Evan’s Blue fluorescence using rainbow dark transformation of Sham, WT, aKO, and nKO groups. Scale bars: 500μm. Three to nine random slices of 3-4 mice per group were included in the data analysis and statistical comparisons (n=16-37). All data are expressed as mean +/- SEM and were compared using a one-way ANOVA with Tukey post-hoc. * indicate significant difference (p<0.05) compared to sham group, and $ indicate significant difference (p<0.05) compared to WT group. Abbreviations: nKO neuronal IGF-1R knockout, aKO astrocyte IGF-1R knockout, WT wildtype, ROI region of interest

Control igfrfl/fl mice (WT) received tamoxifen treatment similar to the inducible knockout mice. Ischemic stroke was induced in the motor and somatosensory cortex via photothrombosis (PTS) one to two months following knockout. Acute changes in neurological response, blood brain barrier integrity, and neuroinflammation were assessed three hours following surgery (Figure 1A). Surgical shams did not exhibit any signs of neurological deficit, while WT PTS mice showed an average neurological deficit score of 2.85±0.26. Comparatively, aKOs had a significantly improved NDS (1.14±0.34) than WTs and nKOs (2.16±0.47) (Figure 1B, F3,25=15.021, n=6-7, p<0.001). No difference in neurological deficits were observed between nKO and WT mice (Figure 1B, p>0.05).

To assess blood brain barrier integrity, mice were administered Evan’s Blue dye immediately following PTS and the distribution of dye was quantified three hours later (Figure 1C-D). We hypothesized that wildtype animals would have increased blood-brain barrier damage compared to sham animals, aKOs, and nKOs. When quantifying the fluorescent Evan’s blue signal within cortical regions subjected to PTS, a main effect of experimental group was observed (Figure 1C, F3,96=11.366, n=16-37, p<0.001). Post-hoc analysis revealed that WT mice exhibited a significant increase in their ROI signal to background (114.33±27.10) compared to shams (9.22±0.626). Interestingly, aKO and nKO mice showed significantly lower levels of Evan’s blue accumulation than WT mice (37.25±3.20 and 44.10±2.52, vs 114.33±27.10), suggesting blood brain barrier leakage was reduced with astrocytic and neuronal IGFR knockout.

### 3.2 Changes in Systemic Inflammation and Microglial Activation in the Hours Following Stroke

Following ischemic stroke, systemic/peripheral inflammation is known to be increased; therefore, we utilized a cytokine/chemokine multiplex panel to measure inflammation in the serum. Table 2 describes the systemic pro and anti-inflammatory production of cytokines/chemokines at three hours following photothrombosis. Of all the cytokines and chemokines of interest, there were only statistical differences between IFN-γ levels between WTs (71.11±11.65) and shams (40.39±1.73) (Table 2). Both knockout models, aKOs (46.77±5.98) and nKOs (54.95±4.42) expressed similar systemic levels of IFN-γ as the sham group (40.39±1.73) (Table 2). No other proinflammatory cytokines or chemokines were significantly upregulated in our mouse model at this early time point.

**Table 2.**
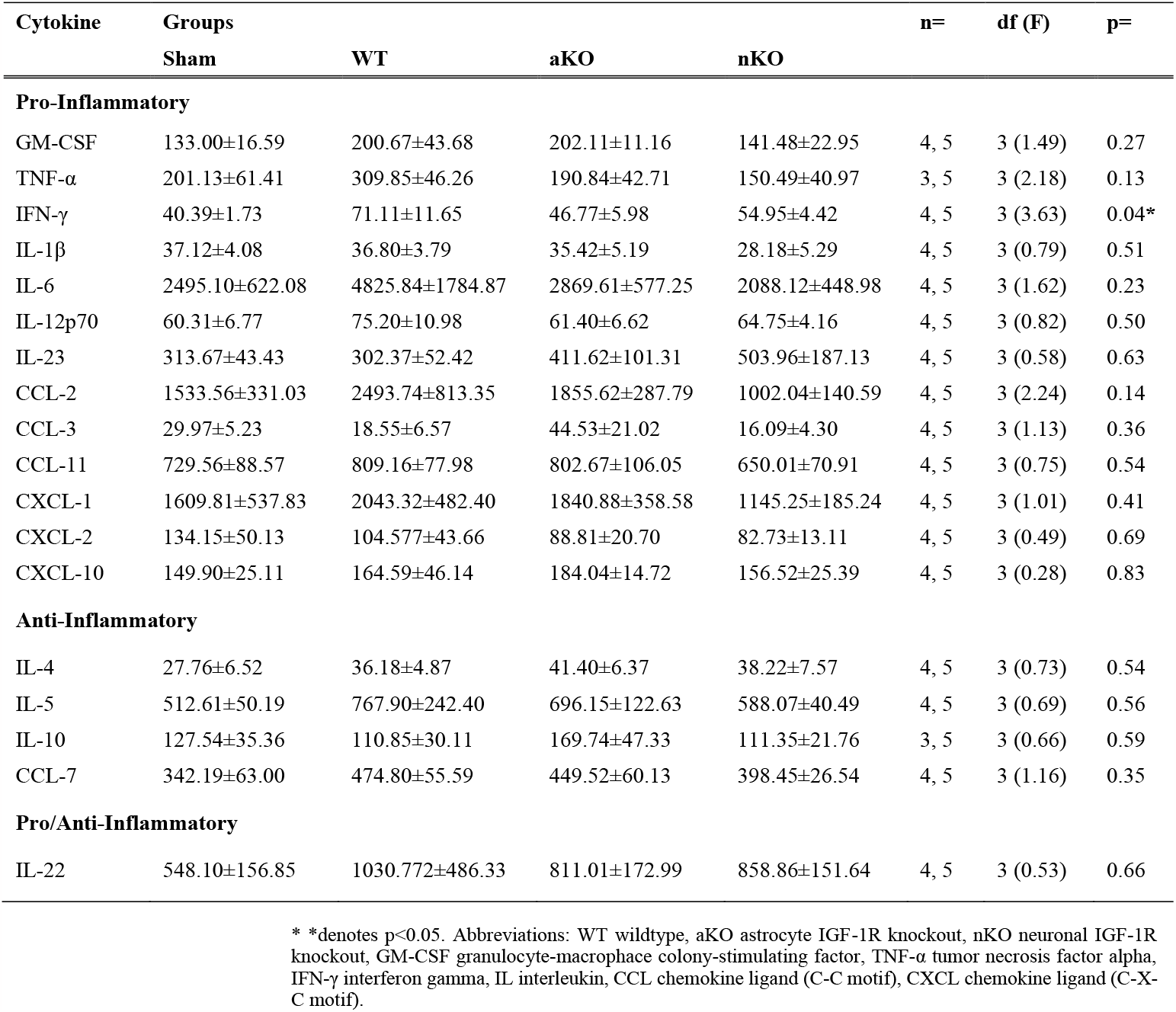
Systemic Inflammatory Profile 3-hours Post Stroke

| Cytokine | Groups |  |  |  | n= | df (F) | p= |
| --- | --- | --- | --- | --- | --- | --- | --- |
|  | Sham | WT | aKO | nKO |  |  |  |
| **Pro-Inflammatory** |  |  |  |  |  |  |  |
| GM-CSF | $133.00 \pm 16.59$ | $200.67 \pm 43.68$ | $202.11 \pm 11.16$ | $141.48 \pm 22.95$ | 4, 5 | 3 (1.49) | 0.27 |
| TNF-α | $201.13 \pm 61.41$ | $309.85 \pm 46.26$ | $190.84 \pm 42.71$ | $150.49 \pm 40.97$ | 3, 5 | 3 (2.18) | 0.13 |
| IFN-γ | $40.39 \pm 1.73$ | $71.11 \pm 11.65$ | $46.77 \pm 5.98$ | $54.95 \pm 4.42$ | 4, 5 | 3 (3.63) | 0.04* |
| IL-1β | $37.12 \pm 4.08$ | $36.80 \pm 3.79$ | $35.42 \pm 5.19$ | $28.18 \pm 5.29$ | 4, 5 | 3 (0.79) | 0.51 |
| IL-6 | $2495.10 \pm 622.08$ | $4825.84 \pm 1784.87$ | $2869.61 \pm 577.25$ | $2088.12 \pm 448.98$ | 4, 5 | 3 (1.62) | 0.23 |
| IL-12p70 | $60.31 \pm 6.77$ | $75.20 \pm 10.98$ | $61.40 \pm 6.62$ | $64.75 \pm 4.16$ | 4, 5 | 3 (0.82) | 0.50 |
| IL-23 | $313.67 \pm 43.43$ | $302.37 \pm 52.42$ | $411.62 \pm 101.31$ | $503.96 \pm 187.13$ | 4, 5 | 3 (0.58) | 0.63 |
| CCL-2 | $1533.56 \pm 331.03$ | $2493.74 \pm 813.35$ | $1855.62 \pm 287.79$ | $1002.04 \pm 140.59$ | 4, 5 | 3 (2.24) | 0.14 |
| CCL-3 | $29.97 \pm 5.23$ | $18.55 \pm 6.57$ | $44.53 \pm 21.02$ | $16.09 \pm 4.30$ | 4, 5 | 3 (1.13) | 0.36 |
| CCL-11 | $729.56 \pm 88.57$ | $809.16 \pm 77.98$ | $802.67 \pm 106.05$ | $650.01 \pm 70.91$ | 4, 5 | 3 (0.75) | 0.54 |
| CXCL-1 | $1609.81 \pm 537.83$ | $2043.32 \pm 482.40$ | $1840.88 \pm 358.58$ | $1145.25 \pm 185.24$ | 4, 5 | 3 (1.01) | 0.41 |
| CXCL-2 | $134.15 \pm 50.13$ | $104.577 \pm 43.66$ | $88.81 \pm 20.70$ | $82.73 \pm 13.11$ | 4, 5 | 3 (0.49) | 0.69 |
| CXCL-10 | $149.90 \pm 25.11$ | $164.59 \pm 46.14$ | $184.04 \pm 14.72$ | $156.52 \pm 25.39$ | 4, 5 | 3 (0.28) | 0.83 |
| **Anti-Inflammatory** |  |  |  |  |  |  |  |
| IL-4 | $27.76 \pm 6.52$ | $36.18 \pm 4.87$ | $41.40 \pm 6.37$ | $38.22 \pm 7.57$ | 4, 5 | 3 (0.73) | 0.54 |
| IL-5 | $512.61 \pm 50.19$ | $767.90 \pm 242.40$ | $696.15 \pm 122.63$ | $588.07 \pm 40.49$ | 4, 5 | 3 (0.69) | 0.56 |
| IL-10 | $127.54 \pm 35.36$ | $110.85 \pm 30.11$ | $169.74 \pm 47.33$ | $111.35 \pm 21.76$ | 3, 5 | 3 (0.66) | 0.59 |
| CCL-7 | $342.19 \pm 63.00$ | $474.80 \pm 55.59$ | $449.52 \pm 60.13$ | $398.45 \pm 26.54$ | 4, 5 | 3 (1.16) | 0.35 |
| **Pro/Anti-Inflammatory** |  |  |  |  |  |  |  |
| IL-22 | $548.10 \pm 156.85$ | $1030.772 \pm 486.33$ | $811.01 \pm 172.99$ | $858.86 \pm 151.64$ | 4, 5 | 3 (0.53) | 0.66 |

\* \*denotes $p<0.05$ . Abbreviations: WT wildtype, aKO astrocyte IGF-1R knockout, nKO neuronal IGF-1R knockout, GM-CSF granulocyte-macrophage colony-stimulating factor, TNF- $\alpha$ tumor necrosis factor alpha, IFN-γ interferon gamma, IL interleukin, CCL chemokine ligand (C-C motif), CXCL chemokine ligand (C-X-C motif).

Microglia have been proposed as an integral modulatory cell type in exacerbating the inflammatory response caused by ischemia and reperfusion. Thus, microglia infiltration and activated phenotypes within the ischemic region were assessed by examining cell count, the number of cellular processes, average branch length, the coverage area, and the average cell size (Figure 2A-F). All main effects of groups are described in the statistical table (Supplemental Materials). Compared to surgical shams, WT mice subjected to PTS showed trending increases in the number of Iba1+ cells (Figure 2B, n=18-24, p=0.073) while both the aKO and nKOs had significantly increased microglial numbers (Figure 2B, n=18-24, p=0.006, p=0.000). Similarly, aKOs and nKOs also had an increase in the percent of area covered by Iba1+ cells (Figure 2C, n=18-24, p<0.001). The elevated count and coverage area within the astrocytic and neuronal IGFR KO mice also corresponded with a statistical increase in the average cell size of both aKO and nKO vs WT (Figure 2D, n=18-24, p<0.001, p=0.024) and shams (Figure 2D, n=18-24, p<0.001), respectively. The average number of microglial endpoints per cell was attenuated in aKO and nKOs (Figure 2F, n=18-24, p=0.049, p=0.001), suggesting a retraction of processes as the cell size grew. Despite this change in process number, the average length of the remaining processes per cell was similar across WT, aKO and nKO mice (Figure 2E, n=18-24, p>0.05).

**Figure 2.**
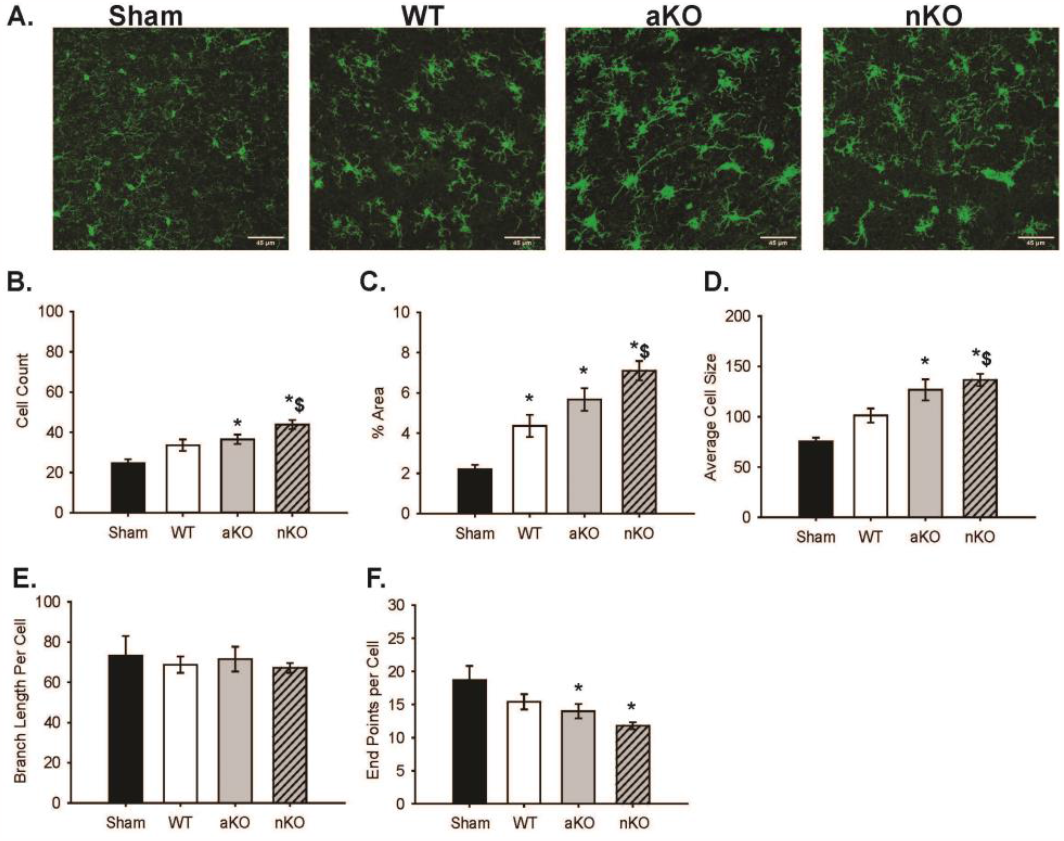
Differences in Microglia Infiltration in the Stroke Cortical Lesioned Area. (A) Maximum z projections of representative images of Sham, WT, aKO, and nKO groups. Scale bars: 45µM. (B) Quantification of IBA1 positive cells, (C) percent area coverage, (D) average IBA1+ cell size, (E) branch length per IBA1+ cell, (F) endpoints per IBA1+ cell, across sham, WT, aKO, and nKO groups. Two random areas of 3-4 brain slices of 3 mice were included in the data analysis and statistical comparisons (n=18-24). All data are expressed as mean +/- SEM and were compared using a one-way ANOVA with Tukey post-hoc. * indicate significant difference (p<0.05) compared to sham group, and $ indicate significant difference (p<0.05) compared to WT group. Abbreviations: nKO neuronal IGF-1R knockout, aKO astrocyte IGF-1R knockout, WT wildtype

### 3.3 Biochemical Changes in the Brains of Cell-Specific IGF-1R Knockouts

Ischemic stroke instigates biochemical changes in proteins, lipids, and other macromolecules that can be detected by Fourier transform infrared resonance spectroscopy (Figure 3) [31, 34, 35]. Representative spectra from WT and astrocytic and neuronal IGFR-KO mice are shown in Figures 3A. The integrated ranges following spectral processing are shown in Figure 3B and Table 1.

**Figure 3.**
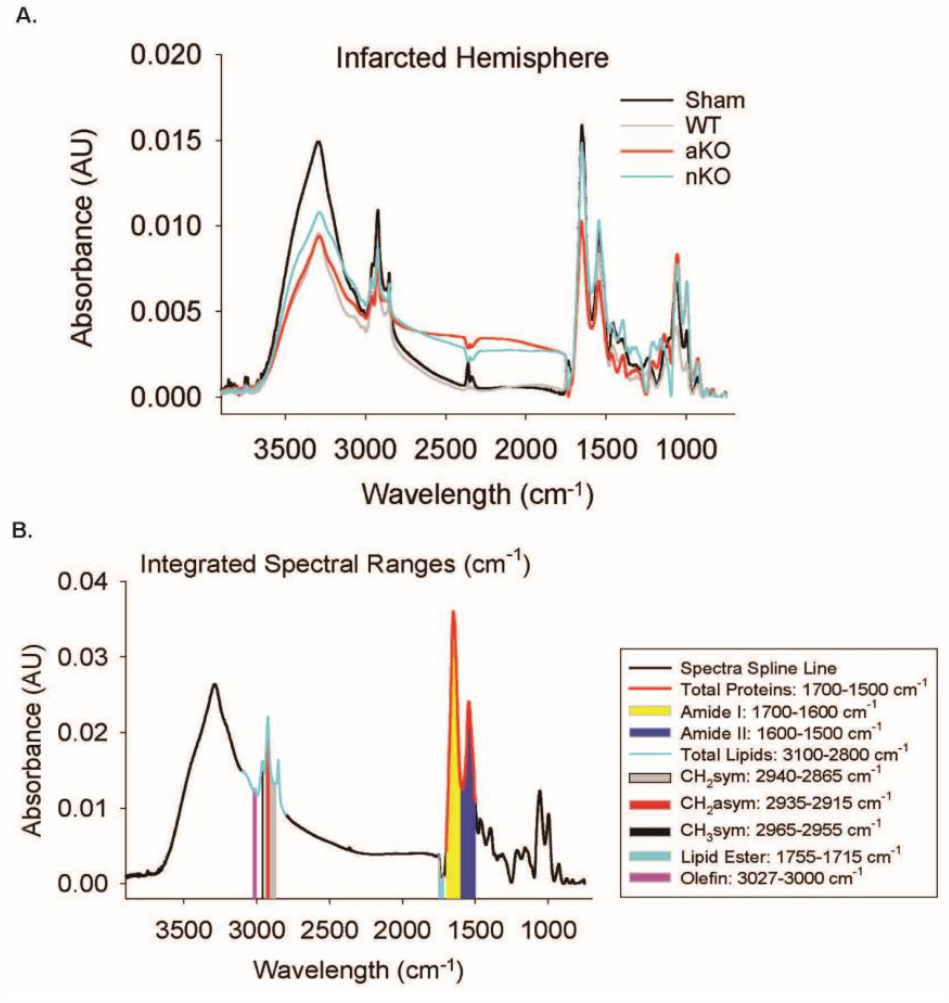
FTIR Spectral Imaging. (A) Representative spectra of the stroke-infarcted hemisphere of Sham, WT, aKO, and nKO groups. (B) Integrated spectral ranges of interest aligned with representative sham spectra. Abbreviations: nKO neuronal IGF-1R knockout, aKO astrocyte IGF-1R knockout, WT wildtype

To determine whether there were baseline differences in chemical content across the transgenic mice, the non-infarcted contralateral hemisphere was assessed (Supplemental Figure 1). There were no observable differences between groups when measuring the Amide I, Amide II, total lipid content, νs(CH2), νa(CH2), and olefin (CH), respectively (n=6, p>0.05). The total protein quantified was significantly higher in nKOs (2.78±0.14) compared to WTs (2.02±0.07) while WTs, aKOs (2.55±0.25), and shams (2.13±0.17) did not differ (F3,20=4.279, n=6, p=0.017), indicating that neuronal IGF-1 reductions increase protein aggregation and manufacturing. The νa(CH3) results were similar, as nKOs (0.003±0.0002) exhibited a higher content compared to WTs (0.002±0.0005) while WTs, aKOs (0.002±0.0002), and shams (0.002±0.0001) did not differ (F3,20=4.801, n=6, p=0.011). On the other hand, WT lipid ester (C=O) content in the contralateral hemisphere (0.002±0.0001) was elevated compared to aKOs (0.005±0.0003) and nKOs (0.001±0.0003) (F3,20=6.761, n=6, p=0.002), suggesting that IGF-1R dysregulation could increase the production of oxidative stress thereby altering lipids. Astrocytic IGFR KOs (0.005±0.0003) and nKOs (0.001±0.0003) had a similar lipid ester (C=O) content as surgical shams (0.001±0.0004) (F3,20=6.761, n=6, p=0.002).

Changes in chemical content were also assessed in the ipsilateral infarcted region 3 hours post-PTS, and numerous changes amongst treatment groups were detected. Unlike in the contralateral side, total protein content did not differ between nKOs (0.77±0.03) and sham mice (1.02±0.01) or WT mice (0.84±0.02); however, aKOs (0.74±0.09) was significantly reduced total protein (Figure 4A, F3,20=3.208, n=6, p=0.045). This astrocyte-specific reduction in total protein following stroke was likely driven by Amide I as, aKOs (0.43±0.05) was reduced compared to shams (0.62±0.04) (Figure 4B, F3,20=5.68, n=6, p=0.005), with no differences being detected with ischemic WT (0.49±0.01) and/or nKO (0.44±0.01) mice. The IAUC of Amide II had similar results; however, WTs (0.17±0.006), aKOs (0.16±0.01), and nKOs (0.16±0.004) all had a lower content compared to the sham surgical group (0.24±0.01) (Figure 4C, F3,20=8.997, n=6, p=0.0005).

**Figure 4.**
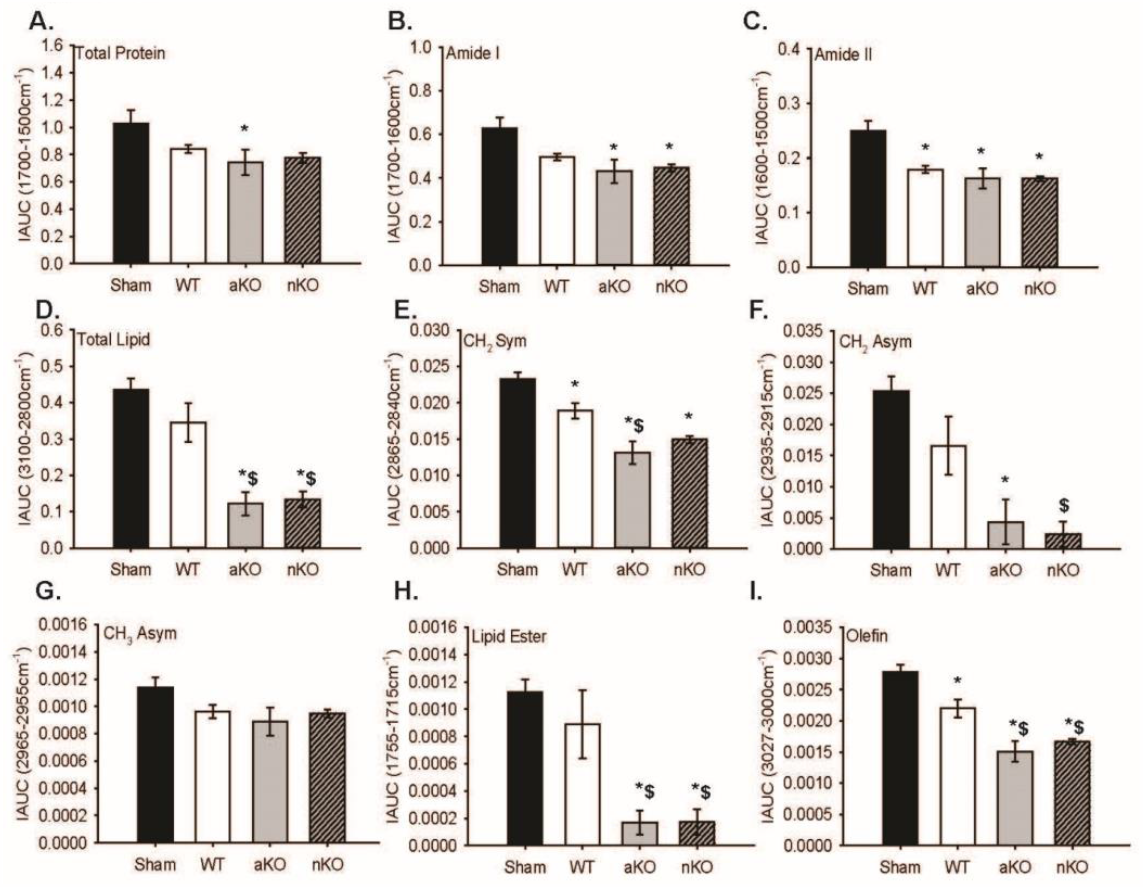
Infarcted Hemisphere Biochemical Changes. (A-I) Quantitative integrated spectral ranges (IAUC) of total protein, Amide I, Amide II, total lipid, νs(CH2), νa(CH2), νa(CH3), lipid ester (C=O), and olefin (CH). Two slices per 3 mice were included in the data analysis and statistical comparisons (n=6). All data are expressed as mean +/- SEM and were compared using a one-way ANOVA with Tukey post-hoc. * indicate significant difference (p<0.05) compared to sham group, and $ indicate significant difference (p<0.05) compared to WT group. Abbreviations: nKO neuronal IGF-1R knockout, aKO astrocyte IGF-1R knockout, WT wildtype

Further analysis revealed that aKOs and nKOs independently had a lower total lipid, νs(CH2), νa(CH2), and lipid ester (C=O) content compared to shams (Figure 4D, F3,20=14.78, n=6, p=2.64E-5; Figure 4E, F3,20=16.77, n=6, p=1.1e-05, Figure 4F, F3,20=10.33, n=6, p=0.0002, Figure 4H, F3,20=11.22, n=6, p=0.0001) indicating lesser tissue lose and oxidative stress.

Astrocytic KOs (0.013±0.001) νs(CH2) amount was lower in WTs (0.01±0.001) as well (Figure 4E, F3,20=16.77, n=6, p=0.00001), while only the nKOs had a lower νa(CH2) content compared to WTs (0.01±0.004). There were no observable differences in the νs(CH3) lipid region (Figure 4G, F3,20=2.44, n=6, p=0.09). Lastly, WTs (0.002±0.0001) had a lower olefin (CH) content that shams (0.002±0.0001), and aKOs (0.001±0.0001) and nKOs (0.001±0.00004) had a lower content compared to shams and WTs (Figure 4I, F3,20=21.04, n=6, p=2.1E-06).

### 3.4 Astrocytic IGF-1R Signaling Influences Neurological Deficits in the Days Following Stroke

Physical changes in the days following stroke were then assessed in a separate cohort of mice. As exogenous IGF-1 has been shown to reduce infarct size and accompanying sensorimotor dysfunction, IGF-1 was intranasally administered to cohorts of WT, astrocytic IGFR-KO and neuronal IGFR-KO mice. One-hour post-stroke, NDS showed main effects of IGF-1 treatment (Figure 5B, F1,96 = 3.938, n=12-20, p=0.050) and genotype (Figure 5B, F1,96 = 5.729, n=12-20, p=0.005). Similar to the three hour timepoint (Figure 1B), at one hour post infarct, WT mice (2.546±0.224) had a significant increase in neurological deficits compared to aKOs (1.542±0.262) (Figure 5B, n=12-20, p=0.012). WT mice also had significantly worse NDS than nKO mice at this time point (1.656±0.212) (Figure 5B, n=12-20, p=0.013). Astrocytic or neuronal KOs did not alter the NDS at one hour (Figure 5B, n=12-20, p=0.939).

**Figure 5:**
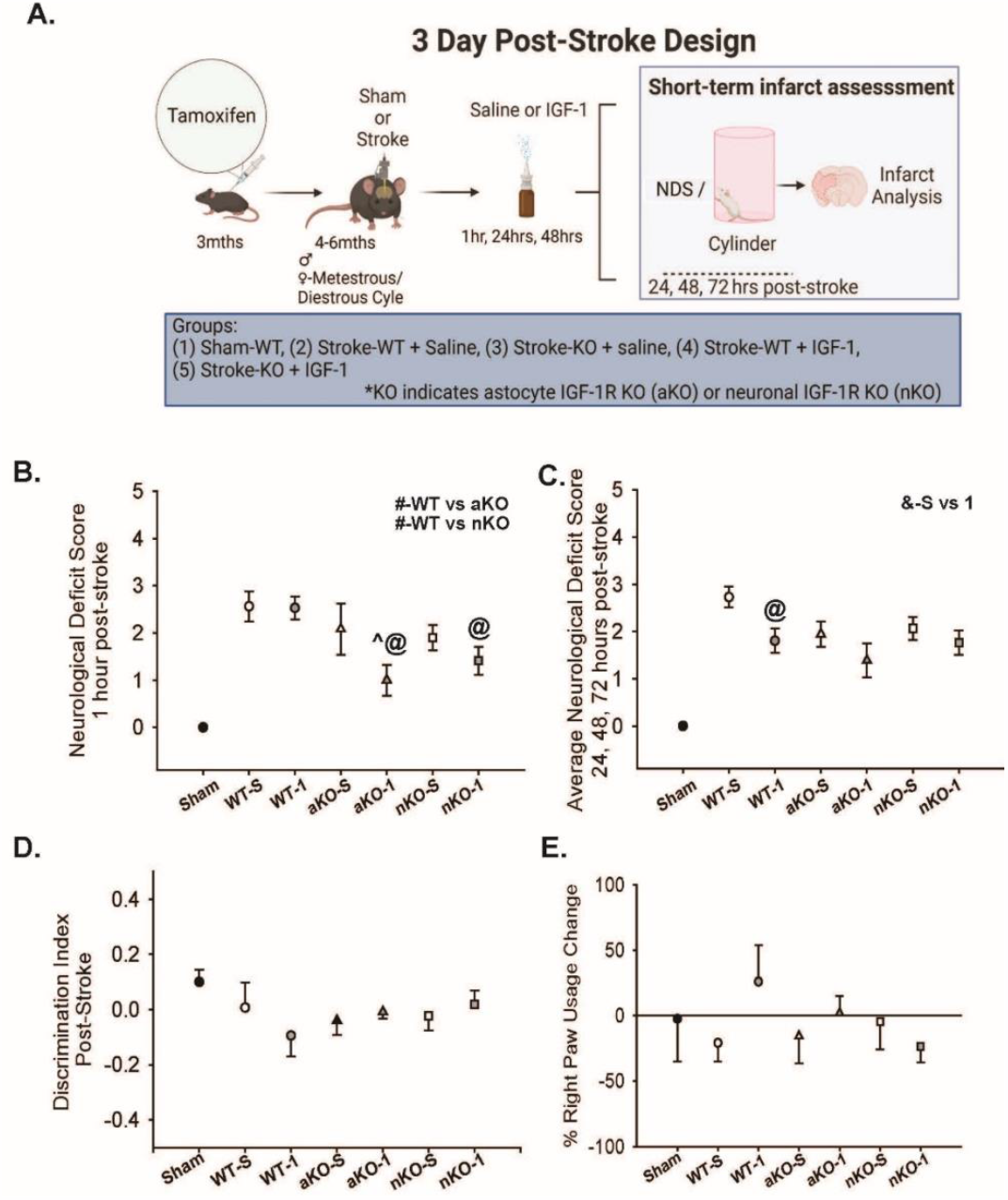
Sensorimotor Coordination Is Not Influence by IGF-1 Signaling in Astrocytes or Neurons. (A) Experimental design and timeline of neuron (nKO) or astrocyte-specific (aKO) deletion of IGF-1R, stroke induction, and behavioral testing. Image made with biorender.com (B) NDS at 1-hour post-stroke (# denotes differences between overall WT/KO groups, ^ indicate significant difference compared to saline treated genotype group, @ indicate significant difference compared to WT-IGF-1 group, p<0.05, n=13-20). (C) Average NDS score of 24, 48, and 72hrs following initial stroke insult (^ indicate significant difference (p<0.05) compared to saline treated group, @ indicate significant difference (p<0.05) compared to WT-IGF-1 group, p<0.05, n=13-20). (D) Discrimination index between right paw and left paw usage 72 hours post-stroke between groups (p>0.05, n=10-19). (E) Percent change in right paw usage 72 hours following stroke NDS (p>0.05, n=6-18). All data are expressed as mean +/- SEM with the error bars beginning from zero to indicate directionality. Sham data points were not included in statistical analysis.Data were compared using a two-way ANOVA with Tukey post-hoc. Equations: discrimination index = ((72hr# - Pre#) / (72hr# + Pre#) + Pre#), % paw usage = ((72hr # - Pre #) / Pre #)*100). Abbreviations: nKO neuronal IGF-1R knockout, aKO astrocyte IGF-1R knockout, WT wildtype, TTC, IGF-1 insulin-like growth factor-1, -S saline treatment, -1 IGF-1 treatment

Groups treated with exogenous IGF-1 (1.647±0.192) exhibited better neurological deficit scores compared to those treated with saline (2.182±0.189) (Figure 5B, F1,96=3.938, n=12-20, p=0.050). Within the IGF-1 treatment, both the aKOs (Figure 5B, n=12-20, p=0.006) and nKOs (Figure 5B, n=12-20, p=0.034) had significantly lower NDS compared to WTs (Figure 5B, n=12-20, p=0.673), suggesting that exogenous IGF-1 provided protection when either neuronal IGFR or astrocytic IGFR was reduced.

When examining interactions between IGF-1 treatment and genotype, IGF-1 x aKO was the only detected interaction (Figure 5B, n=12-20, p=0.042).

Considering the continuous damage in the days following ischemic stroke, we compared NDS across the subsequent 72 hours. An overall difference was observed in the average NDS between saline and IGF-1 treated groups (Figure 5C, F1,96=7.297, n=12-20, p=0.008). In contrastto our results at 1 hour, there was not a main effect difference between genotypes (Figure 5C, F1,96=2.378, n=12-20, p=0.099). Exogenous IGF-1 was sufficient in reducing the averaged NDS of the WTs (Figure 5C, n=12-20, p=0.013), but failed to alter the aKO or nKO genotypes (Figure 5C, n=12-20, p>0.05).

In addition to the NDS, locomotor asymmetry was assessed using the cylinder task. Surprisingly, the mice did not discriminate their paw use towards their contralateral side three days following PTS (Figure 5D, n=10-19, p>0.05). Consistent with the lack of locomotor asymmetry in the discrimination ratio, the percentage of right paw was not statistically different across experimental groups (Figure 5E, n=10-19, two-way ANOVA p>0.05).

### 3.5 IGF-1 Signaling Differentially Preserves Tissue Viability Following Ischemic Stroke

The blood brain barrier permeability, biochemical, and microglial changes observed at the 3-hour time point, as well as the improved NDS scores across the 72 hour observation window led us to hypothesize that astrocyte and neuronal IGF-1R knockouts could exhibit an overall neuroprotective phenotype in the days following ischemic stroke. Main effects of genotype, but not IGF-1 treatment, were observed (Figure 6B) (F1,96=3.811, n=13-20, p=0.026 and F1,96 = 0.607, n=13-20, p=0.607, respectively). Post-hoc analysis revealed trending reductions in aKO (2.674±0.397) infarct damage compared to WT controls (3.853±0.347) (Figure 6B, n=13-20, p=0.071), and significant reductions compared to nKOs (4.030±0.334) (Figure 6B, F1,96 = 3.811,, n=13-20, p=0.026). The infarct size of nKOs (4.030±0.334) and WTs (3.853±0.347) did not differ significantly (Figure 6C, n=13-20, p=0.928).

**Figure 6:**
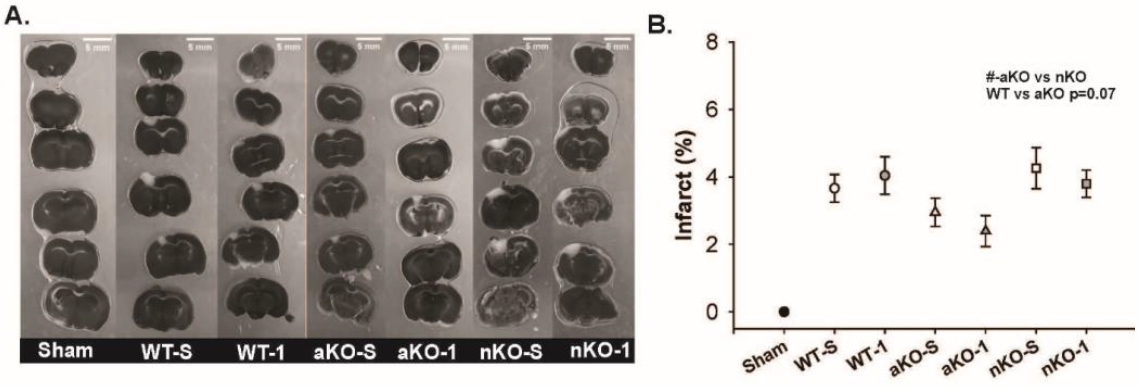
Reductions in IGF-1R Signaling in Astrocytes Attenuates Infarct Size 3-days Post-Stroke. (A) Binary representative images of TTC stained infarct regions (scale bar=5mm). (B) The percent infarct in respective brains regions across saline and IGF-1 supplemented knockouts and wildtype controls. The # indicates a significant difference between aKO and nKO groups (# p<0.05, n=13-20). Sham data points were not included in statistical analysis. All data are expressed as mean +/- SEM and were compared using a two-way ANOVA with Tukey post-hoc. Abbreviations: nKO neuronal IGF-1R knockout, aKO astrocyte IGF-1R knockout, WT wildtype, TTC, IGF-1 insulin-like growth factor-1, NDS neurological deficit score, hrs hours, -S saline treatment, -1 IGF-1 treatment.

## 4. Discussion

To date, effective stroke treatments like tissue plasminogen activator and endovascular thrombectomy have an extremely narrowed therapeutic window, reducing the intervention’s availability. The primary goal of these current therapies are to restore blood flow directly reducing the core of the infarct and indirectly attenuating the exacerbation of the infarct within the penumbral zone. While the infarct core is irreparably damaged, novel therapeutics should be focused on reducing the penumbral damage following initial onset of ischemia. One of those novel therapeutics that has been proposed in IGF-1. Although decades of work have explored the relationship between circulating IGF-1 levels and stroke outcomes, there remains a significant gap in understanding which cells and signaling pathways modulate these findings. In this study, we aimed to compare how cell-specific reductions in IGF-1 signaling in adult astrocytes and neurons influence stroke outcome.

We hypothesized that maintenance of astrocytic IGF-1 signaling would be essential to minimizing tissue damage. The hypothesis was built upon our published in vitro findings that the pharmacological inhibition of IGF-1R in astrocytes impairs glutamate uptake in vitro, reduces glutamate transporter availability, reduces the expression of glutamate handling machinery in vivo, and increases neurotoxicity in triple cell cultures [22]. Moreover, a recent literature review provides comprehensive insights into the essential functions that IGF-1 has in altering glutamate-induced toxicity Ge et al 2022 [36]. This review proposes that IGF-1 signaling could be the primary defense mechanism against glutamate-induced toxicity [36]. In this study, we utilized inducible transgenic mice whereby the IGF-1 receptor could be selectively reduced in adult astrocytes or neurons. We observed strong differences in blood brain barrier permeability, microglia infiltration in the infarct region, and chemical changes within the astrocytic IGF-1R knockout mice in the immediate hours following stroke. However, our aKO mice did not display increased damage; instead, they had less BBB leakage and less neurological deficits. Similar neuroprotective phenotypes were observed with neuron-specific IGFR knock-out mice. These data were surprising as they indicate that IGF-1 signaling on neurons and astrocytes may worsen the immediate changes induced by ischemic stroke.

While BBB leakage was improved with astrocytic and neuronal IGF-1R-knockout, the neuroinflammatory response was not blunted within these transgenic mice. As the resident macrophage of the brain, microglia are responsible for the immune response in the infarct region following ischemic stroke. Interestingly, there was an increase in the number of microglia within the infarcted region of aKO and nKO brains. In fact, nKOs had a significantly higher number of microglia present in the infarct region compared to the aKOs. The increased number of microglia also corresponded to increased cell size, increased microglial coverage area, and retraction of processes-a sign of microglial activation. Thus, our findings indicate that reduced IGF-1 signaling in neurons and astrocytes leads to increased microglial infiltration and activation within the ischemic cortex. While microglia activation leads to increased proinflammatory signaling which could negatively impact neuronal viability and function, the phagocytic activity of activated microglia is also important for cleaning debris and reducing stress within damaged tissue [37]. Therefore, the increased microglial activation within the aKO and nKO mice may contribute to the short-term benefits observed in the NDS and BBB leakage.

We did not quantify proinflammatory cytokines/chemokines within the ischemic tissue but did evaluate systemic inflammation in the hours following stroke as there are known correlations between systemic inflammation and the pro-inflammatory activation of resident microglia. Interestingly, IFN-γ was the only pro or anti-inflammatory that was elevated in the WTs compared to sham, aKOs, and nKOs. The elevations in IFN-γ may lead to a possible increase in the pro-inflammatory subtype of the microglial, however, the lack of change in systemic inflammation in the same animals that showed increased microglial activation could suggest that aKOs and nKOs have a more pronounced restorative microglial activation.

We also assessed histological changes in biochemical contribution in the hours following stroke, as numerous studies have highlighted changes in protein and lipid content within infarcted tissue [38-41]. Ischemia, reperfusion, and ultimately glutamate-induced toxicity is a driving force of oxidative stress that exacerbates stroke damage [42]. Oxidative stress can directly influence protein and lipid damage within ischemic and surrounding tissue. Recent studies have used FTIR imaging to quantify the lipid, protein, and metabolic changes associated with stroke outcomes. In the context of lipids, the production of free radicals induces lipid fragmentation and protein miss-folding in the damaged area which can be observed using FTIR [43]. Of importance, FTIR is the most sensitive and available basic scientific method that can be used to detect lipid, protein, and metabolic biochemical alterations [38-41]. Thus, prior to the onset of necrosis, we utilized FTIR to assess the immediate changes to proteins and lipids that could influence possible neuroprotective phenotypes in our transgenic models. As expected, we did not observe a total protein loss in the ipsilateral hemisphere. These findings are expected considering the narrow (3-hour post-stroke) time window that we assessed these biochemical changes.

Free radical production alters polyunsaturated fatty acid structures (PUFA) inducing lipid fragmentation [44, 45]. Lipid peroxidation can directly increase methyl (CH3) concentrations which are the primary metabolite for additional products [44-46]. Tissue death also induces lipid loss; however, this is primarily affected by necrosis and apoptosis mechanisms, and loss of cells and the integrity of the extracellular environment. From our hypothesis, we did not expect to see differences in in total lipid or protein content, or in lipid products: lipid acyl (CH2) and lipid methyl (CH3); moreover, the treatment with PFA serves to diminish the detectable lipid content in tissue with FTIR. Nevertheless, our findings show that ischemic stroke aKOs and nKOs exhibit lower total protein and lipid content which emphasizes the possibility that a greater tissue loss could proceed over time, as well as an increase in oxidative stress that could alter biochemical degradation. Specifically, we observed a significant loss of lipid ester (C=O) and olefin (CH) content in the ischemic stroke aKOs and nKOs. Our biochemical assessment using FTIR showed that the loss of IGF-1 signaling in astrocytes and neurons exacerbated total lipid loss and lipid peroxidation production. Thus, it was not surprising that neuronal IGFR-knockout mice did not show improvements in infarct size three days later. We assume that the increased lipid damage in the astrocytic knockout mice would also have led to a failure of neuroprotection three days later. However, it is possible that a protective mechanism was triggered in the aKOs at the immediate time of insult that allowed them to overcome this macromolecular damage. This thought is supported by the improved NDS scores in aKO mice in the immediate hours following ischemic stroke.

A recent meta-analysis revealed that lower IGF-1 levels were overall associated with reduced stroke risk [47]. Moreover, several previous studies have shown that exogenous IGF-1 administered to rodent models of ischemic stroke reduces infarct size and accompanied functional deficits [48-53, 16, 26]. We hypothesized that exogenous IGF-1 would reduce the negative outcomes of ischemic stroke in WT mice, and that this protection would be mediated through either astrocytes or neurons so one of the transgenic mutants would fail to show this protection (aKO or nKO). We observed partial protection with IGF-1, whereby IGF-1 treated groups exhibited fewer neurological deficits when compared to the saline treated groups. We also observed a reduction in the NDS in the IGF-1 treated aKOs compared to the saline treated aKOs, indicating that IGF-1 was capable of exerting beneficial effects independent of astrocytic IGFR. Neurons mayhave an increased availability of free IGF-1 in the aKO model facilitating the presence of the neuroprotective phenotype with a reduced infarct size. If local production of IGF-1 were increased in aKO mice, then the observed resilience to ischemic damage in the aKO mice would be expected. This direction of thinking could also corroborate why the improved neurological responses appear to be time restricted, as over time (24-72 hours) these IGF-1 levels deplete, leaving neurons vulnerable to death per usual. The aforementioned possible mechanism of immediate protection and then circumvention to tissue death is a biphasic one that needs further investigation.

Our results indicated that intranasal IGF-1 was insufficient in reducing the infarct size. Considering it affected the neurological deficit scores, we do believe that it was bioavailable when delivered in this route. Previous studies using intranasal administration of IGF-1 in stroke studies were conducted in adult and neonatal rats along with the MCAO or hypoxic-ischemia method which provides reasoning for why our treatment was insufficient to induce neuroprotection [26, 55, 56]. Although intranasal administration of drugs is a promising technique to bypass the BBB, this technique produces variability in mice across research groups. Many previous intranasal studies use various methods, but the most common is the use of anesthesia to consistently administer a certain number of microliters of drug over time. This method has significant implications in stroke studies considering anesthesia is neuroprotective against ischemic stroke [28]. To combat this, we administered a smaller volume of IGF-1 to awake mice which could indicate why there were not infarct changes. Moreover, intranasal IGF-1 may be a more targeted method for ischemia and reperfusion methods like the transient middle cerebral artery occlusion (MCAO) model but considering PTS does not induce a significant penumbral size, we could intuitively expect it not to be a sufficient therapeutic for this rodent model. Ultimately, we chose PTS over MCAO, due to its highly reproducible method to induce ischemic infarcts in an extremely controlled region.

We did not observe any differences in the locomotor asymmetry in the cylinder task when calculating the discrimination index and the ipsilateral/contralateral paw usage post stroke. Yet, there are apparent directional changes in the use of paws. One possible reason that we did not see a significant change is the variation seen in the parameters of individuals that have utilized the cylinder task. For example, the majority of our mice did not reach a total of 20 paw touches during the 5-minute duration of the task along with us habituating our mice with a preliminary test prior to inducing stroke. These minor deviations in the protocol could explain the lack of observed differences, while additional tests like the Catwalk could be a more precise study to measure locomotor changes.

## 5. Conclusions

In conclusion, we observed that ischemic stroke aKOs and nKOs exhibited reduced blood brain barrier damage but increased number and size of microglia within these groups. In addition, we observed significant genotypical changes in the biochemical assays that are influenced by ischemic stroke. Exogenous IGF-1 administration continued to provide benefits in both neuronal IGF-1R KO mice and astrocytic IGF-1R KO mice, indicating that neither cell type was the primary mechanism by which IGF-1 exerts neuroprotection. Double knock-outs or additional knock-outs within the other cell types that contribute to neurovascular structure/function should be considered. Importantly, the beneficial effects of IGF-1 signaling manipulations appear to be temporally restricted as damage continued to accumulate in both knock-outs over a 3 day observation window. Together, these findings provide a significant insight into how cell-specific neuroendocrine signaling mechanisms are crucial to disease onset and expansion of damage following ischemic stroke.

---

**Supplementary Table:**
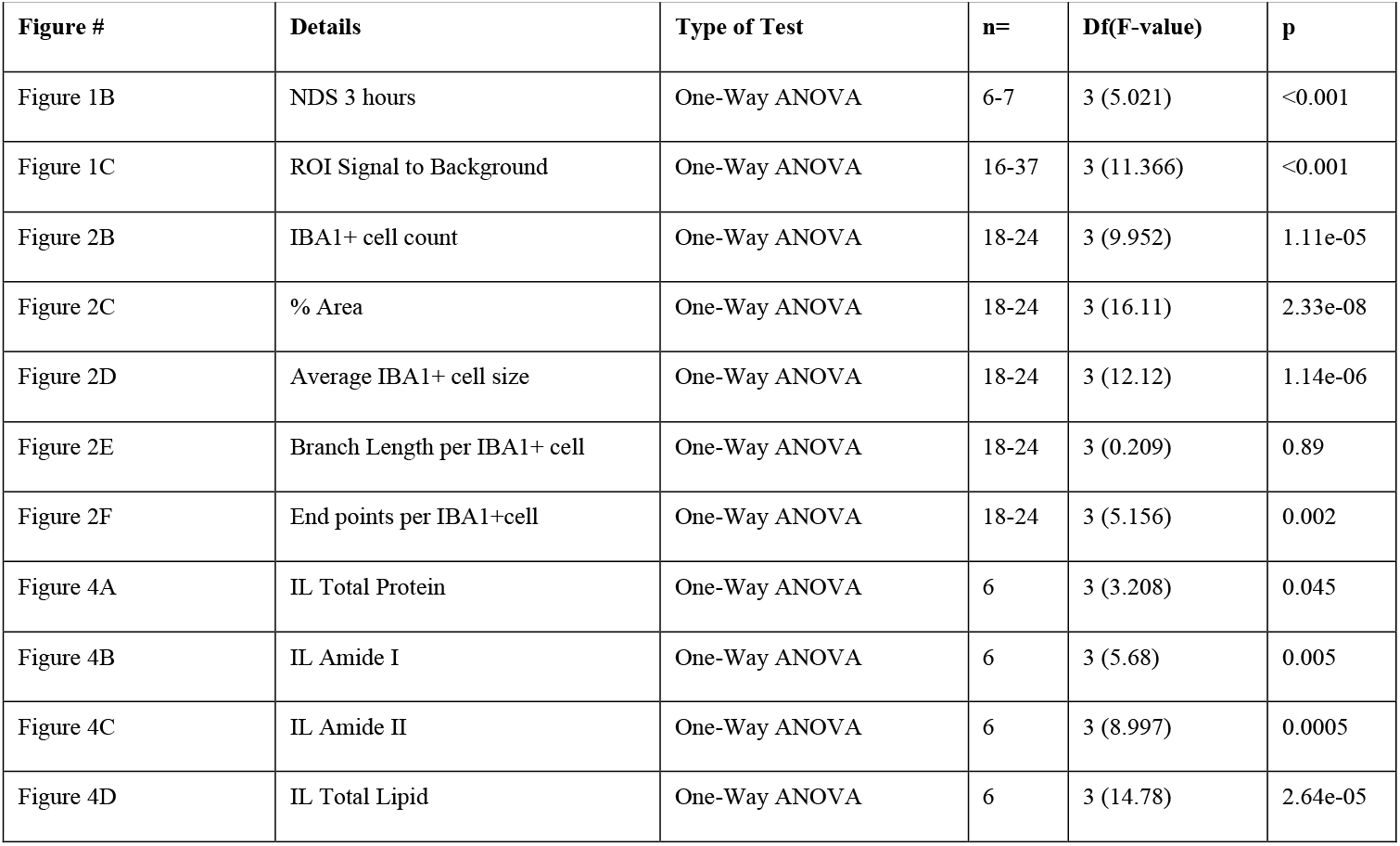
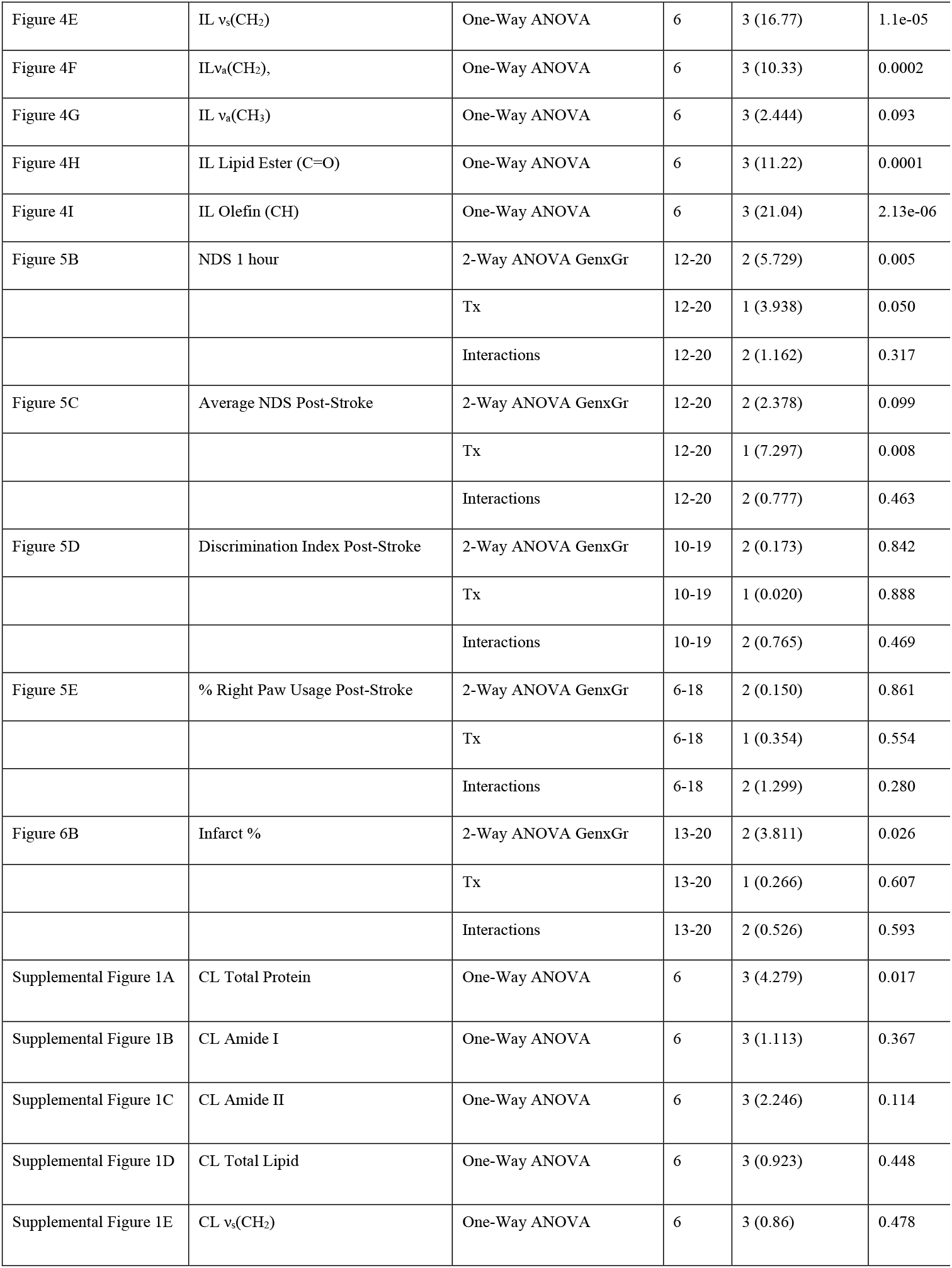
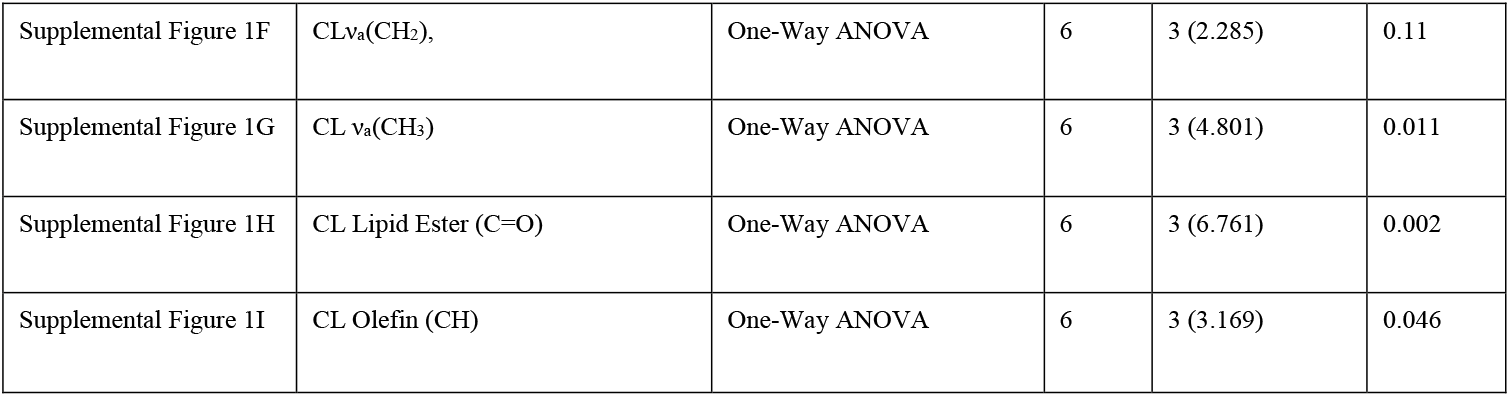
Statistical Details.

**Supplemental Figure:**
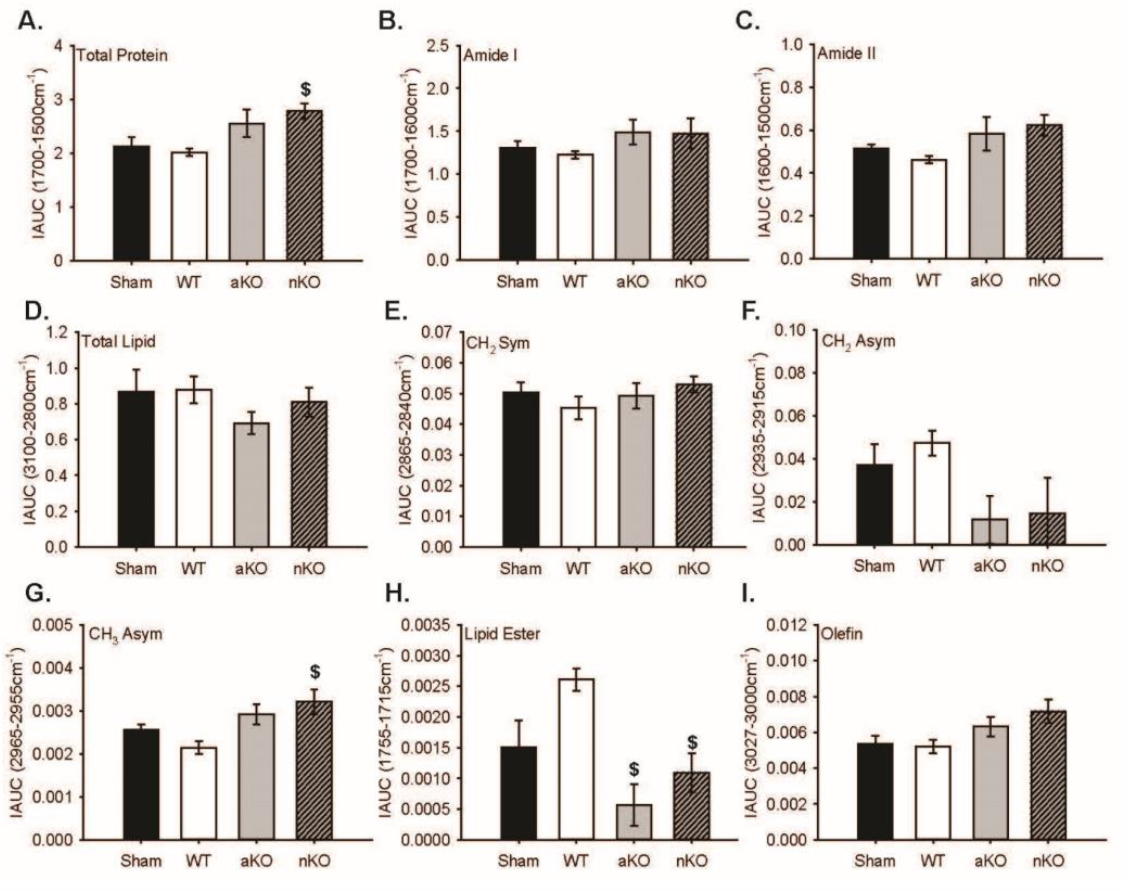
Healthy Hemisphere Biochemical Changes. (A-I) Quantitative integrated spectral ranges (IAUC) of total protein, Amide I, Amide II, total lipid, νs(CH2), νa(CH2), νa(CH3), lipid ester (C=O), and olefin (CH). Two slices per 3 mice were included in the data analysis and statistical comparisons (n=6). All data are expressed as mean +/- SEM and were compared using a one-way ANOVA with Tukey post-hoc. * indicate significant difference (p<0.05) compared to sham group, and $ indicate significant difference (p<0.05) compared to WT group. Abbreviations: nKO neuronal IGF-1R knockout, aKO astrocyte IGF-1R knockout, WT wildtype

## Author Contributions

CH and NA were responsible for conceptualization, investigation, project administration, and formal statistical analyses. KW contributed to acquiring the FTIR images. MP assisted with formal analysis of FTIR images. MCH, and NA developed and designed methodology. BA, AV, KT, and NM performed genotyping, cylinder acquisition, blinded cylinder analysis, and curated data.MD assisted with Tamoxifen injection. CH prepared visualizations. CH and NA prepared the original draft, and all authors reviewed and edited the manuscript.

## Acknowledgments

The authors would like to thank the animal care staff at the University of Mississippi.

